# DeepCryoPicker: Fully Automated Deep Neural Network for Single Protein Particle Picking in cryo-EM

**DOI:** 10.1101/763839

**Authors:** Adil Al-Azzawi, Anes Ouadou, Max R Highsmith, John J. Tanner, Ye Duan, Jianlin Cheng

## Abstract

Cryo-electron microscopy (Cryo-EM) is widely used in the determination of the three-dimensional (3D) structures of macromolecules. Particle picking from 2D micrographs remains a challenging early step in the Cryo-EM pipeline due to the diversity of particle shapes and the extremely low signal-to-noise ratio (SNR) of micrographs. Because of these issues, significant human intervention is often required to generate a high-quality set of particles for input to the downstream structure determination steps. Here we propose a fully automated approach (DeepCryoPicker) for single particle picking based on deep learning. It first uses automated unsupervised learning to generate particle training datasets. Then it trains a deep neural network to classify particles automatically. Results indicate that the DeepCryoPicker compares favorably with semi-automated methods such as DeepEM, DeepPicker and RELION, with the significant advantage of not requiring human intervention.

---

To build a reliable high-resolution 3D reconstruction of protein structures from Cryo-EM images, one must extract hundreds of thousands of single particle images from 2D cryo-electron microscopy [1] [2]. The use of high-energy electrons can result in radiation damage to specimens during imaging and result in extremely noisy micrographs, and consequently a limited electron dose is preferred [3] [4]. The signal-to-noise-ratio (SNR) of original (2D) micrographs tends to be very low, with noise from a variety of sources including low contrast, particle overlap, ice contamination and amorphous carbon [5]. Hence, the task of single particle picking is still challenging in some cases [5]. Many different computational methods have been proposed for the semi-automated single particle picking over the past decades. Single particle picking using template-based matching methods are very sensitive to noise [6] [7] [8] [9] [10] [11] [12]. Thus, some initial “good references” have to be selected in advance to ensure that those manually selected examples have less noise compared with the other in the same (2D) micrographs. Similarly, the edge-based [13] [14] and feature-based methods [15] [16] [17] show significant reduction in performance since they are sensitive to the lower contrast of the (2D) micrographs [5]. Deep learning methods for single particle picking have been proposed, including EMAN2.21 [18], DeepEM [5], DeepPicker [19], and FasetParticlePicker [20]. These deep learning methods made significant contributions to addressing the particle picking issue. However, there are some unsolved challenges such as lack of diversified training dataset, high false positive rate, and the difficulty of dealing with low-SNR micrographs.

Over the past decade many different computational methods have been proposed for the automated and semi-automated single particle picking tasks. These methods are based on different techniques such as template-based matching, edge detection, feature extraction, and conversional computational vision [3]. Recently, Deep Learning has exponentially grown in the field of machine learning. Many Deep Learning algorithms from the field of computer vision use convolutional techniques to extract features from big data via layers in neural networks [11]. Furthermore, deep learning appears to be a suitable approach for cryo-EM image processing as the size and number of the micrographs per data set are continually increasing while the SNR of micrographs remains low [3]. Several deep learning approaches to particle picking have been proposed, including EMAN2.21 (particle picking with convolutional neural network [18]), DeepEM [5], DeepPicker [19], and FasetParticlePicker [20].

Hence, we propose a fully automated deep neural network for single particle picking based on the fully automated training particle data generation using unsupervised learning algorithms. We use two clustering approaches (regular clustering algorithm using the Intensity-Based Clustering IBC) [21] and super clustering algorithms using the super k-means [22]) to automatically generate training particle datasets for training the deep neural networks. To accommodate the low-SNR cryo-EM images, a general framework of micrograph preprocessing that has been used in both our last two models [21] [22] is applied to improve the quality of the low-SNR micrographs.

The method is tested on cryo-EM images of the Keyhole Limpet Hemocyanin (KLH) [23], Apoferritin [24], 80S ribosome [25], and β-galactosidase [26]. A key feature of our approach is the use of Non-Maximum Suppression (NMS) [27] during the testing phase in order to reduce the number of false-positive particle detections. Overall, the automated DeepCryoPicker improves the performance of particle picking over semi-automated methods such as DeepEM, DeepPicker and RELION-2 (using referenced-based picking) [28].

## Results

### Overview of the DeepCryoPicker procedure

DeepCryoPicker is designed for fully automated single particle picking in cryo-EM. Our framework contains two components: The first component is a training particle-selection algorithm based on unsupervised learning (shown on the left side of Fig. 1(a)). The second component is single particle picking utilizing supervised deep learning (shown on the right side of Fig. 1(a)). The first component has two sections: automated training particles picking, and automated training dataset generation. The first section of the automated training particles selection is based on two steps. Firstly, the micrograph images are pre-processed using a set of advanced image processing tools to enhance and increase the quality of the micrographs. Secondly, each cryo-EM image is clustered using two different unsupervised learning clustering algorithms and then each clustered image is cleaned and used to detect and isolate each particle. Some irrelevant objects are removed. The second section of the automated training particle selection is based on automatically evaluating each isolated particle sample and classifying it as either a “good” or “bad” training sample. The second component is the fully automated single particle picking method based on a deep learning scheme which has two steps. The first step is designing and training a deep convolutional neural network using the training dataset that has been automatically generated using the first component of our framework. In the second step, the trained model is used to test every micrograph after pre-processing them using the same preprocessing stage that is used to prepare the training dataset. Two different micrograph testing datasets are used for testing.

**Fig. 1.**
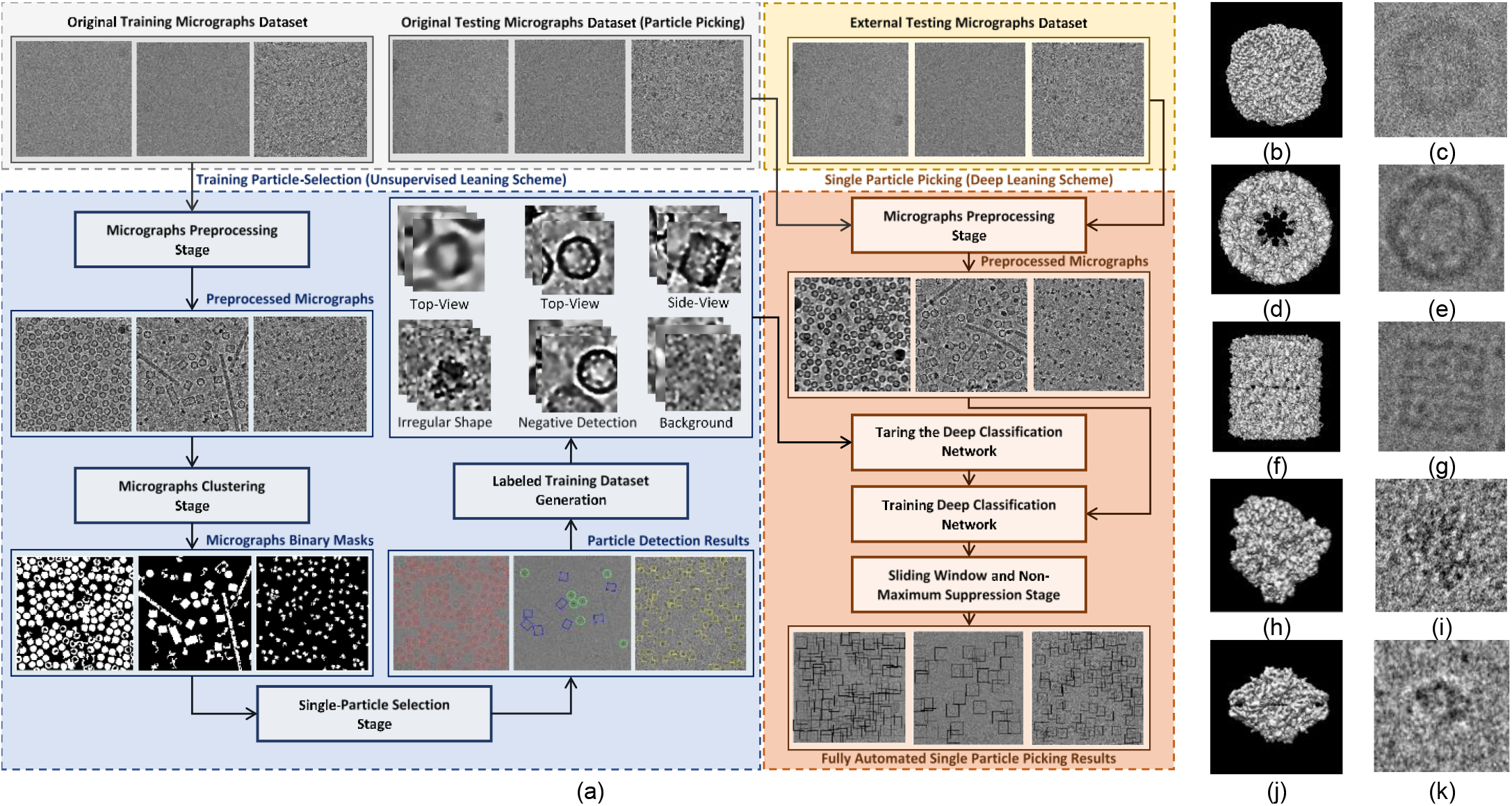
(a) The general workflow of the training particle-selection based unsupervised scheme and single particle picking based on deep learning scheme. The gray part of the workflow shows the micrographs data collection. The blue part of the workflow shows the fully automated training particles-selection using clustering algorithms. The red part of the workflow shows the general flow of the single particle picking using deep classification network. The yellow part of the workflow shows the external testing part of the DeepCryoPicker, (b) 3D Cryo-EM map of apoferritin, (c) picked particle from an apoferritin micrograph, (d) 3D Cryo-EM map of KLH viewed from the top, (e) picked particle from a KLH micrograph showing the top view (circular particle), (f) 3D Cryo-EM map of KLH viewed from the side, (g) picked particle from a KLH micrograph showing the side-view (square particle), (h) 3D Cryo-EM map of the 80S ribosome, (i) picked particle from a ribosome micrograph, (j) 3D Cryo-EM map of beta-galactosidase, (k) picked particle from a beta-galactosidase micrograph.

### Micrographs data collection

We consider three typical protein shapes in micrographs that are collected from different micrograph datasets as shown in Fig.1. The first protein shape is circular, as exemplified by the apoferritin [24]. Its 3D Cryo-EM map is shown in Fig. 1(b), while a picked particle is shown in Fig.1 (c). The second protein shape is square, as seen in the side-view of KLH [23]. There are two main types of projection views in this dataset. The top view is circular (Fig. 1(d-e)), while the side view is square (Fig. 1(f-g)). The third protein shape that is considered is the general case of an irregularly-shaped protein such as the 80S ribosome (Fig. 1(h-i)) [25] and β-galactosidase (Fig. 1(j-k)) [26].

### Experiments on unsupervised learning framework for fully automated training particles-selection

The automated training particle selection model has two steps: automated training particle picking, and automated training dataset generation. In the first step, 80% of the samples from the collected micrographs are used. Numerous particles are composed and picked from micrograph images using the fully automated framework for particle picking based unsupervised learning approaches that we proposed in our previous models [21] [22]. Then, each single particle image is automatically isolated and evaluated as a “good” or “bad” training sample. The total number of particles for each dataset is shown in Table.1.

**Table 1.** Total number of the training particles-selection using fully automated good training particles-selection for apoferritin, KLH, and Ribosome datasets.

| Criteria | Apoferritin Top-View | KLH Top-View | KLH Side-View | Ribosome Side-View |
| --- | --- | --- | --- | --- |
| Total Particle Image Picking | 356 | 878 | 356 | 1276 |
| Particle Size | 178x178 | 221x221 | 221x225 | 187x187 |
| Perfect Particle Circle Shape Selection | 337 | 874 | 337 | 1157 |
| Non-perfect Particle Shape Selection | 19 | 4 | 19 | 19 |
| Number of Images | 20 | 82 |  | 20 |
| Size of Micrograph | 1240x1240 | 2048x2048 |  | 4096x4096 |

### Experiments on automated training dataset generation

The final training dataset has five classes. Three classes that represent the original particle shapes (top-view, side-view, and irregular (complex) protein shapes) are automatically selected from the “good” particle examples after evaluating every single particle. The image samples of the other two classes are automatically generated from different micrograph’s background as “background class” or automatically expanded and collected from the “bad” training samples as “negative detection class”. Then, a certain number of image samples are randomly selected from each training class to expand the size of the training dataset. A sample is rotated 90, 180, and 270 degrees to generate three additional training samples. The total number of training particles before and after regeneration are shown in Table.2.

**Table 2.** Automated training particles-selection datasets. The second column illustrates the total of the particles picked from the training micrographs before applying the good training particles selection and automated training dataset generation and expansion, while the third column illustrates the total number of particles after applying the good training particles selection and automated training dataset generation and expansion.

| Dataset | Before Re-generation | After Re-generation |
| --- | --- | --- |
| Apoferritin | 921 | 1500 |
| Ribosome | 1157 | 1500 |
| KLH (Top-View) | 874 | 1500 |
| KLH (Side-View) | 337 | 1500 |
| Negative detection | - | 1500 |
| Background | - | 1500 |

### Experiments on training deep learning classification models

To understand the impact of the number of classes on the classification model, we varied the number of classes in the training dataset via three different experiments. In the first experiment, we used all five classes to train and validate the deep classification model. In the second experiment we remove the “background” class while keeping the other four classes. In the third experiment, we remove the “negative detection” class while keeping the other four classes. The corresponding precision-recall curve of each experiment in the training dataset showing that in the third case (using three main classes and background class), yields the best result with an average precision of 100%. The average precision is reduced to 98% and 99% in the first and second case respectively.

### Experiments on testing deep learning classification models

To evaluate the three deep learning models above, we split our dataset into training, testing and validation sets. Each class has 1500 particle images, we split the data to 80% for training and validation (1200 particle images, 960 for training and 240 for validation) and 20% testing (300 particle images). The total number of the training particles using 5 classes in the first case is 5250 particles while the total number of the testing particles is 2250. For the second and third models with either background class or negative class, the training set contains 4200 particle images and the testing set contains 1800 particle images. The error or loss of the deep neural network was used as a feedback parameter to tune and adjust the weight and bias, including the number of the feature maps, kernel size of the convolutional layers, and the subsampling kernel size of the subsampling layer. Moreover, the training/testing cycles were tuned based on the hyper-parameters and updated the training datasets until the accuracy of the deep neural network reached a satisfactory level. Fig.2 shows some testing examples of the deep classification network after training based on the third experiment type (three main particle shape classes and background class). The testing accuracy of the deep classification networks using different number of classes in Table 3. It is clear that the deep classification model achieves a higher accuracy 99.89% based on using the three particle classes plus the background cases.

**Fig. 2.**
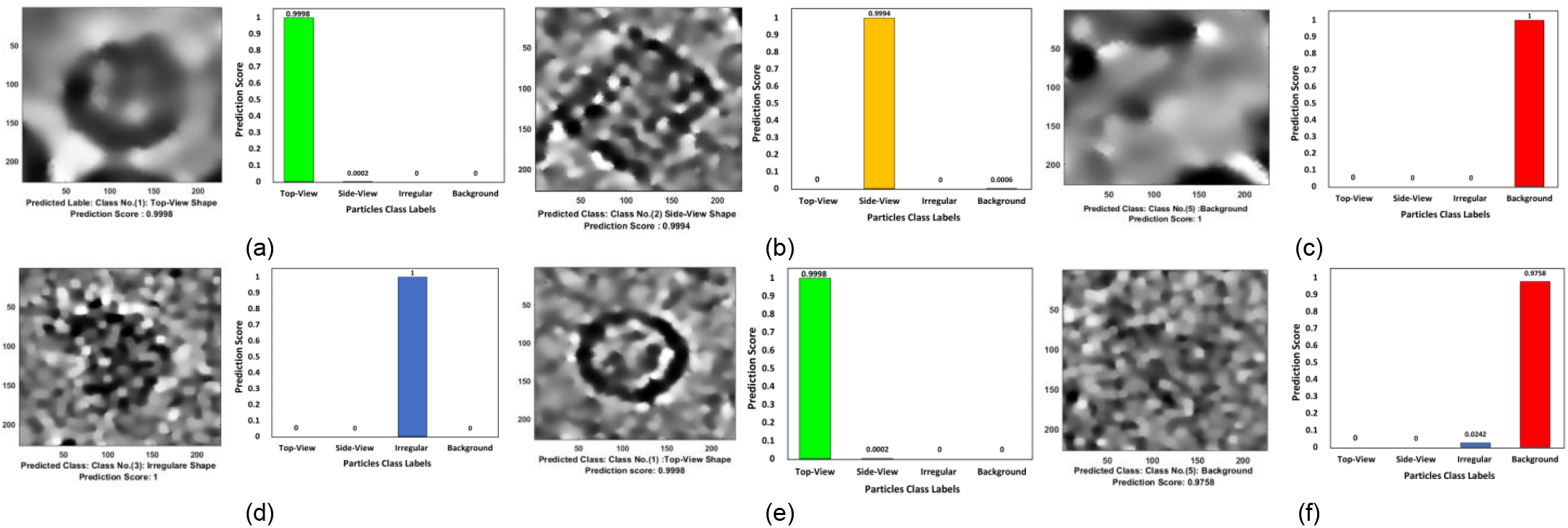
Different examples of the deep classification network results. (a) A typical testing image example showing high-density top-view particle’s predicted label and prediction score of the apoferritin micrograph dataset, (b) A typical testing image example showing high-density side-view particle’s predicted label and prediction score of the KLH micrograph dataset, (c) A typical testing image example showing high-density background predicted label and prediction score, (d) A typical testing image example showing high-density irregular particle’s predicted label and prediction score, (e) A typical testing image example showing high-density top-view particle’s predicted label and prediction score of the KLH micrograph dataset, (f) A typical testing image example showing high-density background predicted label and prediction score.

**Table 3.**
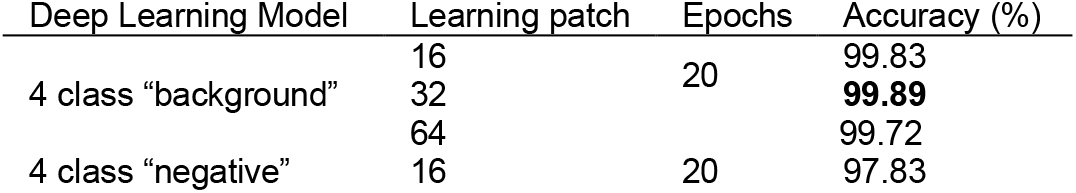
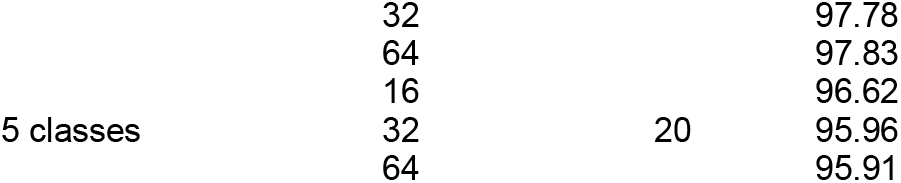
Test results of using different parameters and datasets. Epochs: number of iterations of training.

### Experiments of deep learning framework for fully single particle picking on cryo-EM datasets

The second component of our DeepCryoPicker is the fully automated single particle picking. It has three steps: scanning test, scoring cleaning, and filtering using non-maximum suppression. In the first step, a sliding-testing window is used to scan each micrograph in the testing dataset from the top left to the bottom right corner with a constant step size. To determine the prediction parameter, a fixed size sliding window (square box) is chosen to be slightly larger than the particle size. During the scanning-testing step, each single patch is extracted and fed to the trained deep classification network. Each sliding window receives a certain prediction value [0 1] from the deep network model. The prediction scores represent the probability there is a particle at the center of the corresponding window. In the second step, a scoring map is generated for each tested micrograph. The scoring map describes the likelihood score distribution of the particles over the entire micrograph. In fact, some detected object such as ice or noise can be predicted as a particle (i.e. false positive). To discard the false positive detection, a cleaning step is implemented which connects any two pixels in scoring maps whose prediction scores are close and both above the threshold. Then, a connected area (pixels) is regarded as a false positive if the size of connected area is larger than a cutoff value and is removed from the candidate list. Finally, we use non-maximum suppression (NMS) [27] to refine the current particle candidate list. NMS is used to filter the detection boxes based on their Intersection over Union (IoU) between the detected boxes. The candidate particle filtering based on the NMS has three main steps: sorting, selecting, and repeating. First, all candidates’ boxes for each given particle category are sorted based on their prediction scores (from high to low). Second, the candidate box that has the highest prediction score is selected as the final candidate box. Then, all other candidate boxes within the selected IoU are discarded. Third, among the remaining boxes the NMS repeats the two second steps until there is no remaining box in the candidate list.

A typical result of DeepCryoPicker is shown in Fig.3 and Table 4, which illustrates the results of the particle picking using the fully automated framework and different micrographs from different datasets. The average precision-recall both reached 97%. Fig.4 shows the precision-recall curves for each particle shapes individually using different datasets such as apoferritin, KLH [23] (the top-view particle shapes), KLH [23] (only the side-view particle shapes), Ribosome and β-galactosidase (irregular and complex particle shapes). For instance, Fig.4a shows the blue plotted curve of the precession-recall for top-view particle shapes picking, Fig.4b shows the red plotted curve of the precession-recall for side-view particle shapes picking, and Fig.4c shows the black plotted curve of the precession-recall for irregular and complex particle shapes picking.

**Fig. 3.**
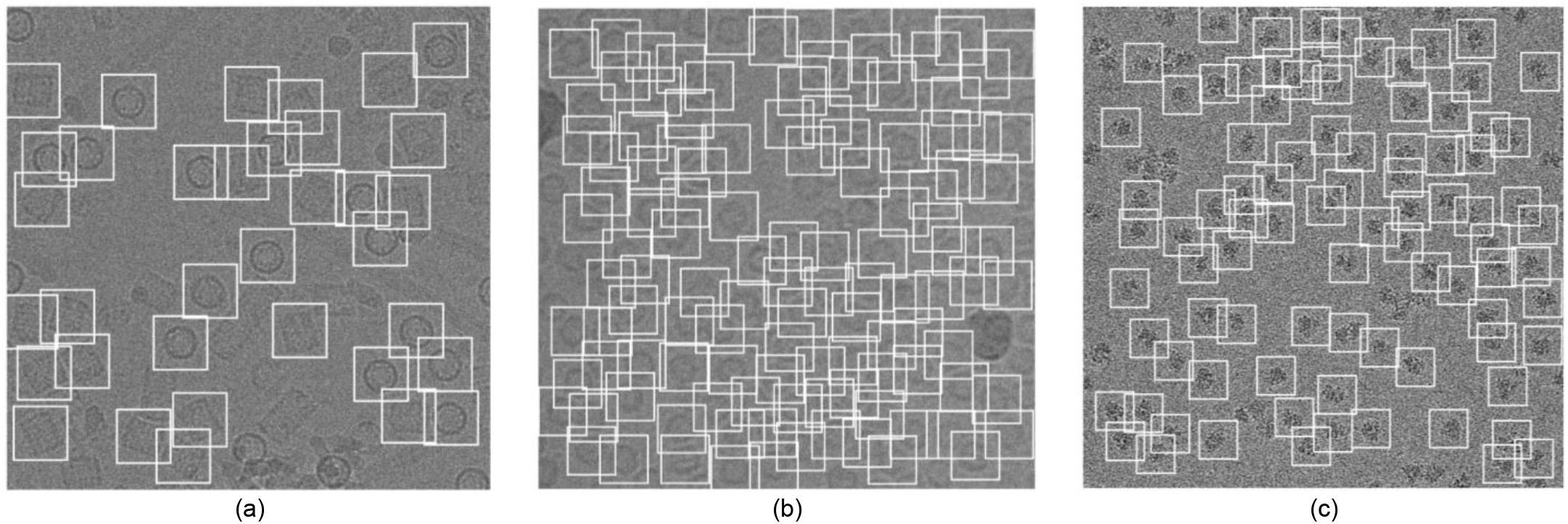
The results of DeepCryoPicker on three micrographs. (a) Top and Side-View particles picking on an image in the KLH dataset, (b) top-view particle picking on an micrograph in the apoferritin dataset, (c) irregular (complex) particle picking on a micrograph in the Ribosome dataset,

**Fig. 4.**
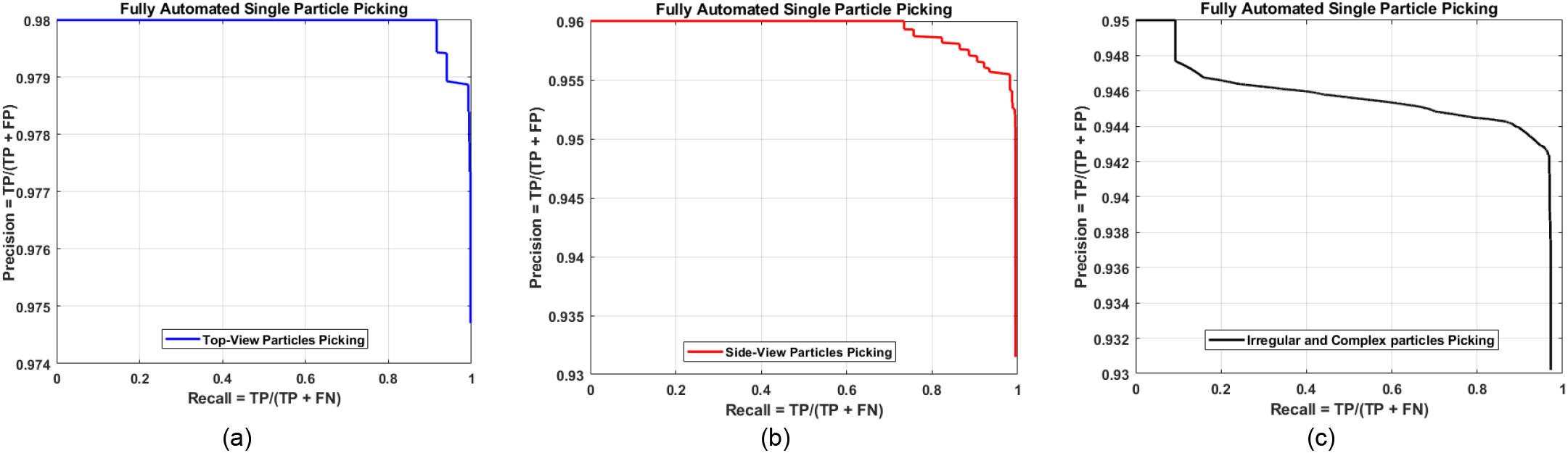
Precision-recall cures of the fully automated different single particle shapes picking result using deep classification network and different micrographs datasets, (a) precision-recall cure of the top-view particle shapes picking, (b) precision-recall cure of the side-view particle shapes picking, (c) precision-recall cure of the irregular and complex particle shape picking.

**Table 4.** DeepCryoPicker Evaluation table using different micrograph datasets. Our model is trained for all views (top-view, side-view, and irregular particles) in additional to two optional classes (Background and Negative samples).

| Trained For | Accuracy | Precision | Recall | F1 Score |
| --- | --- | --- | --- | --- |
| 4 classes with Background | 0.97 | 0.94 | 0.98 | 0.96 |
| 4 classes with Negative | 0.96 | 0.99 | 0.92 | 0.96 |
| 5 classes with Background and Negative | 0.95 | 0.95 | 0.88 | 0.92 |

### Experiments on unseen testing micrographs datasets

In addition to testing our model on different test micrographs (testing sets) split from the whole datasets and in terms of the generalization our model to unseen datasets, we further test our model on three different micrographs (external testing micrographs) of other proteins that are different from those of training and test datasets (Fig. 5). The external testing micrographs have been selected based on different particle shapes (Fig. 5a, b and c). For instance, Fig. 5a is an external testing micrograph from the bacteriophage MS2 (EMPIAR-10075) [29] where the particle shapes are identical top-view. Fig. 5b, shows another external testing result on an external testing micrograph from the T. acidophilum 20 (EMPIAR-10186) [30] where the particle shapes are either top-view or side-view. Finally, Fig. 5c, shows the last external testing result on an external testing micrograph from beta-galactosidase 2.2 Å EMPIAR-10061) [31] where the particle shapes are irregular shapes.

**Fig. 5.**
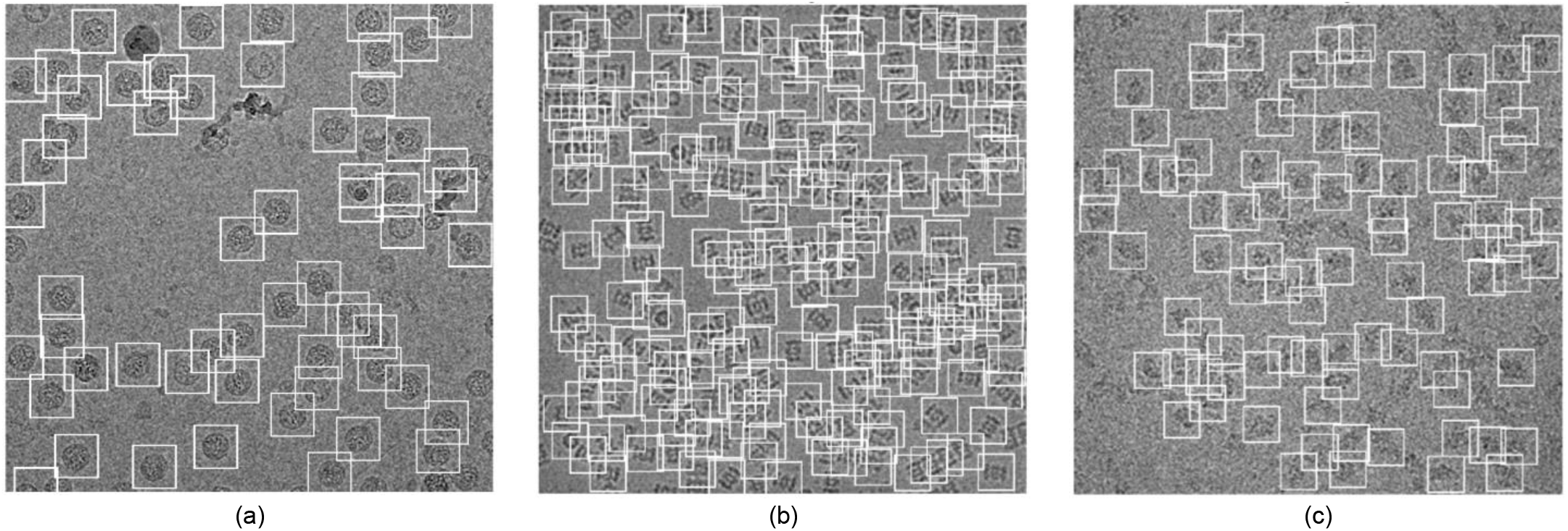
The DeepCryoPicker testing results using different micrographs from different external testing datasets (a) typical external micrograph from the bacteriophage MS2 (EMPIAR-10075) [28] showing the Top-View particles picking, (b) typical external micrograph from the T. acidophilum 20 (EMPIAR-10186) [19] showing the Top and Side-View particles picking, (c) typical external micrograph from the beta-galactosidase 2.2 *A*°(EMPIAR-10061) [31] showing the irregular (complex) particles picking.

### Computational efficiency

We used a desktop computer equipped with an NVIDIA GeForce GTX 1070 graphics card GPU with 4 GB memory and an Intel Core i7 6900K CPU to train DeepCryoPicker. The time needed for training was 22–144 min for the whole dataset (Fig. 6).

**Fig. 6.**
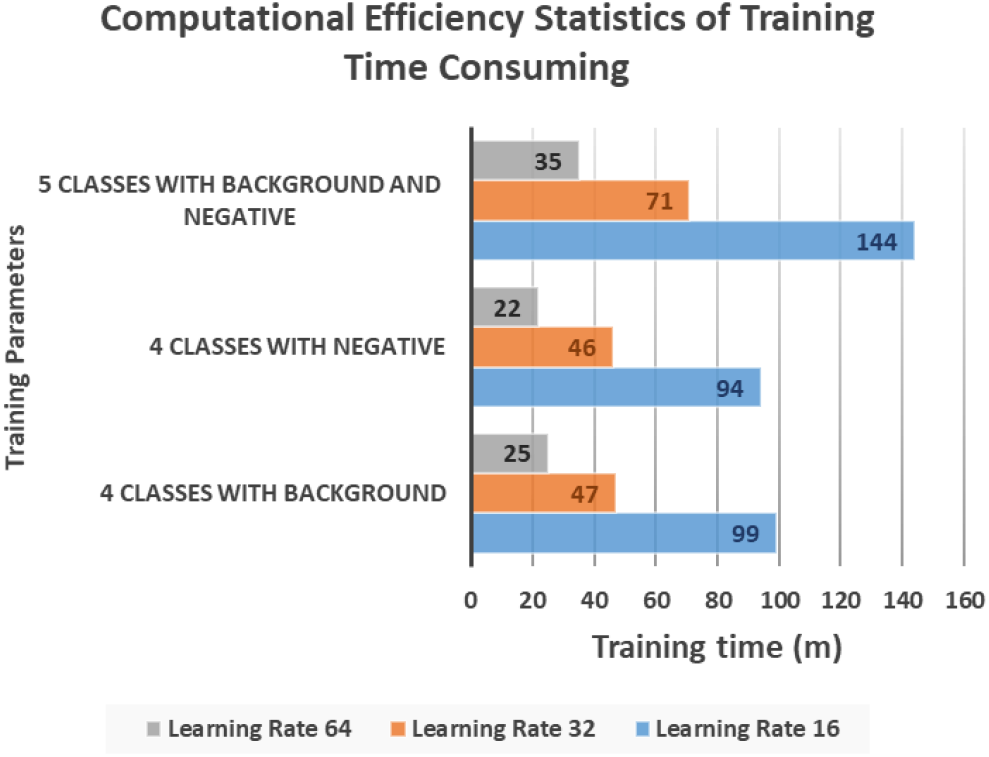
Computational efficiency statistics of DeepCryoPicker training times.

## Discussion

Our method tackles significant challenges that other particle picking approaches have faced such as lack of diversified training dataset, high false positive rate, and the difficulty of dealing with low-SNR micrographs. First, to generate a sufficiently large training dataset, we design a fully automated training particle selection based on the unsupervised learning algorithms. Most of the regular protein shapes (circles and squares) have been fully automated picked based on our IBC algorithm. And most of the irregular and complex protein shapes have been accurately picked based on a fully automated unsupervised learning approach using the super k-means clustering algorithm. Therefore, the generation of the training set is fully automated, eliminating the need for manual labeling or labor-intensive particle selection. Second, to accommodate the low-SNR images, a general framework of micrographs preprocessing [21] [22] is applied to improve the quality of the low-SNR micrographs. In general, the preprocessing steps increase the particle’s intensity and pre-grouping the pixels inside each particle makes them easier to be isolated. Third, to reduce the number of false-positive (FP) particle detections, we use Non-Maximum Suppression (NMS) [27] during the testing phase. It removes duplicates of bounding boxes centered around the same region, consequently decreasing false-positive detections.

We compare the results from the DeepCryoPicker with different particle picking tools such as RELION-2 [28], PIXER [3], DeepPicker [19], and DeepEM [5] using the KLH [23] datasets. Fig. 7 shows the precision-recall curves of these methods. The green, yellow, black, blue, and red curves represent the precision-recall curves for RELION-2 [28], DeepPicker [19], DeepEM [5], PIXER [3], and DeepCryoPicker respectively. The results indicate that DeepCryoPicker performance is slightly better than RELION-2 [28], with the advantage of being fully automated. The improvement of DeepCryoPicker’s performance over other methods is more pronounced.

**Fig. 7.**
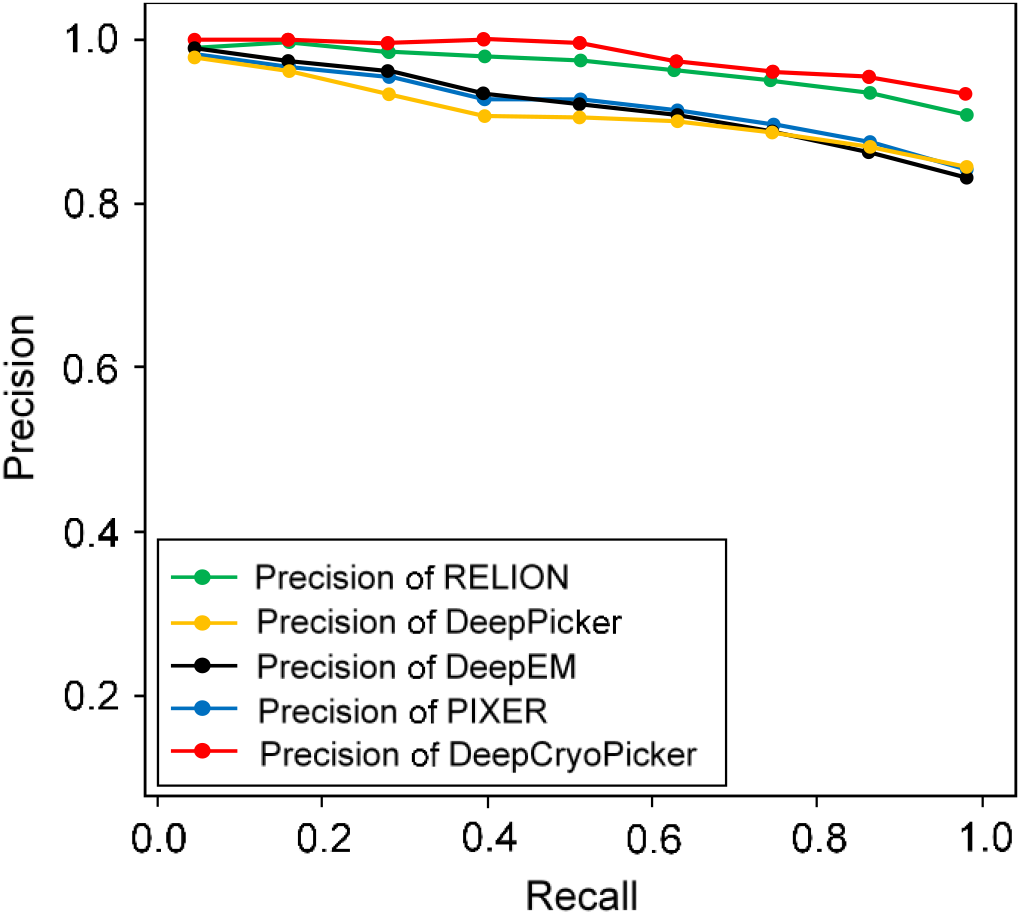
The precision-recall curves of particle picking for different single particle picking tools. The green, yellow, black, blue, and red curves represent the precision-recall curves for RELION-2, DeepPicker, DeepEM, PIXER, and DeepCryoPicker respectively.

## Methods

DeepCryoPicker consists of two components (Fig.7): (1) Component 1: fully automated training particles-selection based on unsupervised learning; (2) Component 2: fully automated single particle picking based on deep classification network. The orange rectangle marks the first part of the fully automated approach “fully training particles-section and dataset generation” while the dark blue rectangle marks the second part “fully automated single particles picking”. The green and gray rectangles mark the first and second stage of the preprocessing step. The blue boxes at the top denotes the datasets used in this work.

### Component 1: fully automated training particle selection based unsupervised learning approach

This component consists of two stages: (1) Stage 1: fully automated training particle selection; (2) Stage 2: fully automated perfect “good” training particle selection and labelled training dataset generation.

### Stage 1: Fully automated single particle-picking

Two different fully automated single particle picking approaches based on unsupervised learning (AutoCryoPicker [21] and SuperCryoPicker [22]), are used in this stage. AutoCryoPicker [21] and SuperCryoPicker [22] used the same preprocessing procedures to increase the SNR and the quality of each micrograph as shown in Fig. 8 (green and gray rectangles). The results of the preprocessing procedures for apoferritin [24], KLH [23], Ribosome [25], and Beta-galactosidase [26] images are shown in Fig. S1 (in the Supplementary File). The particle picking results are shown in Fig. S2 and Fig. S3.

**Fig. 8.**
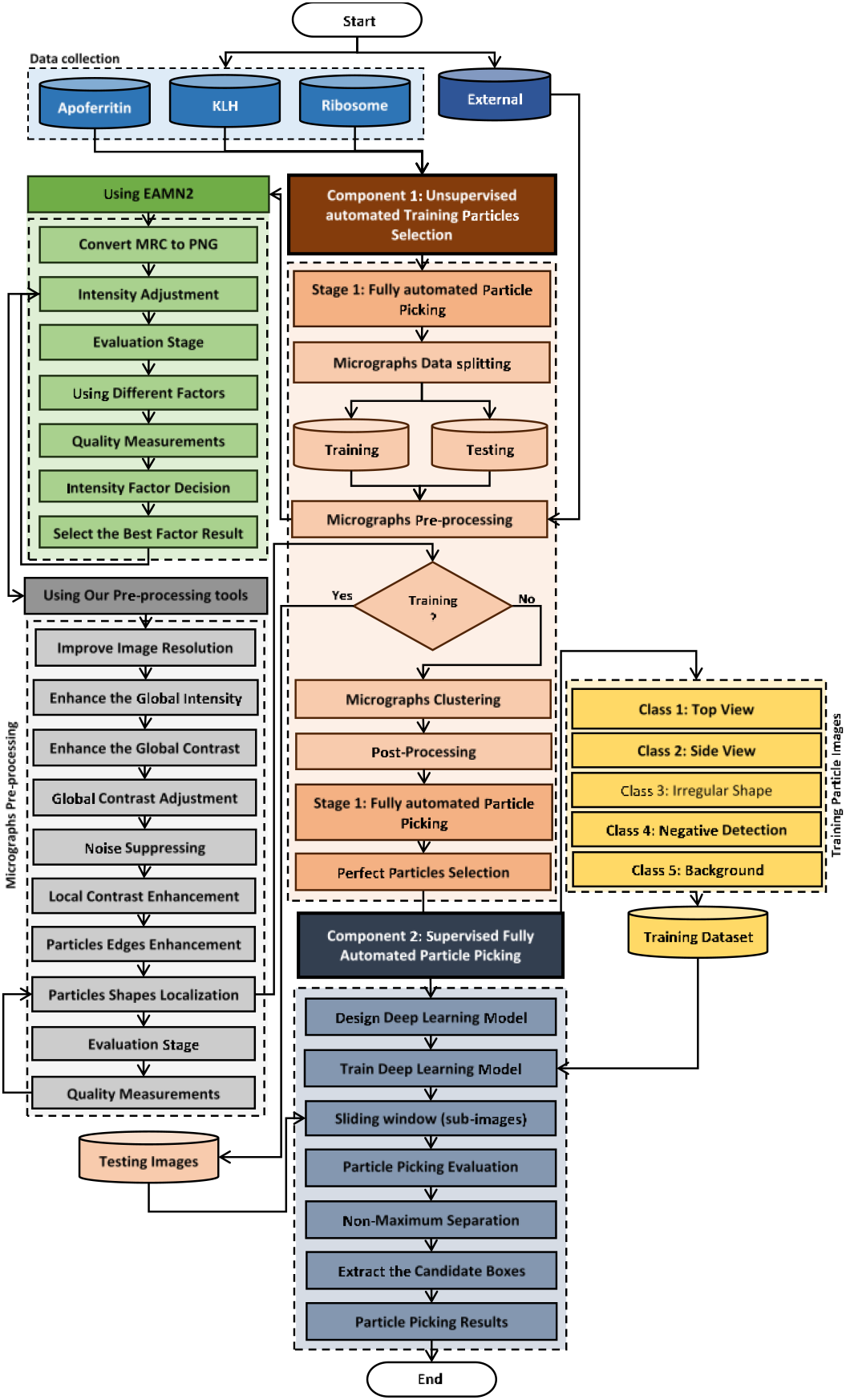
DeepCryoPicker workflow. The orange rectangle marks the first part of the fully automated approach “fully training particles-section and dataset generation”. The blue rectangle marks the second part “fully automated single particles picking”. The green and gray rectangles mark the first and second stage of the preprocessing step respectively.

### Stage 2: Fully automated training particles-selection

In this stage, each single particle that comes from the first stage is evaluated and selected as a good training example using three fully automated perfect “good” training particles-selection approaches.

### Perfect “good” top-view training particle selection

We develop an additional step called “good top-view (circular) training particle selection” (see Algorithm in the Supplementary File). This step is based on using the individual binary mask for each particle as shown in Fig.S4d, f, h, and j (in the Supplementary File). We use the modified Circular Hough Transform algorithm (CHT) in AutoCryoPicker [21] to generate a perfect circle on top of each particle’s mask. Then we test each individual particle’s mask size and verify if it is a perfect full circle and label it as either a “good example” or a “bad example”. We test each top-view particle by calculating the average roundness value for the whole top-view (circular) particles. This is determined by computing the area and perimeters using the connected component particle mask’s pixel index list and the circularity based on the Equation (1):

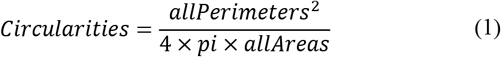

where *allAreas* is the area of each selected particle and *allPerimeters* is the cemetery size of each particle. Each individual particle (circular) that achieves the average object roundness class is considered as a “good” training example, otherwise as a “bad” training example. Fig. S5 (in the Supplementary File) shows the results of the good top-view training particle selection. Fig. S5a and S5e show individual top-view particle binary masks from the apoferritin [24] and KLH [23] datasets. It is noticed that a perfect circle has been successfully drawn on top of the particle’s binary mask using the modified CHT algorithm as shown in Fig. S5b and f. Fig. S5c and g show the replaced artificial perfect circle binary masks that will be used later to test the particles for apoferritin [24] and KLH [23] datasets. Fig. S5d and h show the good apoferritin [24] and KLH [23] top-view training particles selection. In contrast, Fig. S5i, l, m, and o show other examples of the top-view particle’s binary masks that the modified CHT has failed to draw perfect circles on top of them. Fig. S5j, l, n, and p show some bad top-view training particle examples.

### Perfect “good” side-view training particles-selection

For the side-view particles picking, we do not have issue with the overlapped particle selection since the only prefect side-view (square) particles are selected through the side view (square) training particle shape selection in cryo-EM based on using the “overlapped particles removal and perfect side-View particles selection algorithm” in the AutoCryoPicker model [21]. Fig. S6(a) and (g) in the supplemental document show different KLH cryo-EM clustering results using Intensity-Based Clustering Algorithm (ICB). Fig. S6(b) and (h) show the KLH cryo-EM clustered images after the circular and non-square object removal. The binary mask images have only the square particle shapes (side view) in the whole cryo-EM images. Some overlapped particles still exist in the cleaned binary mask as is shown in Fig. S6(b). The overlapped particles are removed from the final cleaned masks (See Fig. S6e and f) after applying the overlapped particles removal using Feret diameter measures approach [32] (see Fig. S6d and j). Fig. S6f and l show the same KLH binary mask images after the perfect side-view (square) particles shape generation on the of the cleaned binary masks. Fig. S7 and S8 in the supplemental document show an example of the perfect side-view (square) particle selection. Fig. S8a and (d) show different cryo-EM micrographs form the KLH dataset. Fig. S8b and e show the final results of side-view particles-selection using different micrographs form the KLH dataset based ICB clustering, and perfect square (side view) particle shapes detection using Feret object diameter [32]. Fig. S8c and f show also the top-view particles-selection results based on modified ICB clustering, and modified CHT [21].

### Perfect “good” irregular and complex training particle selection

This step is also based on using the individual binary mask for each complex and irregular particle as shown in Fig. S9b, d, f, and h. Then, we test each individual particle’s mask size and determine if it is a usable training sample. We develop a “good irregular (complex) training particle selection” algorithm (see Algorithm S2 in the supplemental document) to test each irregular binary particle, by calculating the average area for the whole particle binary masks which is determined by computing the total number of white pixels in each particle using the connected component particle mask’s pixel index list. Then the average area as is shown in Equation (2):

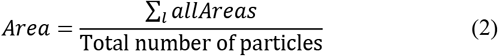

where *l* is the total number of particles in each cryo-EM image.

### Component 2: fully automated single particle picking based on deep classification network

The second component of the DeepCryoPicker is the particle picking based deep network shown in Fig.9. It consists of many layers such as input layer, pre-processing layer, convolutional layers, sub-sampling layers, two fully connected layers, and one output layer. The main architecture of the DeepCryoPicker has in total thirteen layers as is summarized in Table 5. The first and second layers (input and the pre-processing layer) come from the first component of the DeepCryoPicker. The input layer takes the particles that have been already picked through the first model of the DeepCryoPicker. Each particle has been picked based on the preprocessed version of each of the micrographs. The rest are five convolutional layers, three max-pooling (subsampling) layers, two fully connected layers, and one output layer. To use one deep network structure, we unify the variety of the particle sizes as shown in Table 1 to one fixed size. In this case, after each particle is detected, a bounding box is drawn around each particle object in the cryo-EM image which is used to crop the particle image from the original micrograph. We recalculate the bounding box dimension of each detected particle after calculating the center of each box and specifying the fixed size of each (width and height). Then, the input size of the first and second layer (input and the preprocessing) in our DeepCryoPicker structure is 277 × 277. The third layer is the convolutional layer using 96 kernels with size 11 × 11. the first convolutional layer (third layer in the structure) produces 96 feature maps with dimensions 55 × 55. The fourth layer is the max-pooling layer with kernel size 3 × 3 and the feature maps output dimension is 27 × 27. The fifth layer is another convolutional layer using 256 kernels with size 5 × 5. The fifth layer (convolutional) produces 256 feature maps with dimensions 27 × 27. The sixth layer is another max-pooling layer with kernel size 3 × 3 and the feature maps dimensions output is 13 × 13. The seventh, eighth, and ninth layers are convolutional layers using different numbers of kernels 384, 384, and 256 respectfully. We use the same kernel size 3 × 3 for three convolutional layers. The output feature maps size for the last three convolutional layers 13 × 13. The tenth layer is the third max-pooling later with kernel size 3 × 3 and output dimensions 6 × 6. The last two layers are the fully connected layers to the final output (prediction layer) where the particle class is predicted based on the weight’s matrix and the activation function.

**Fig. 9.**
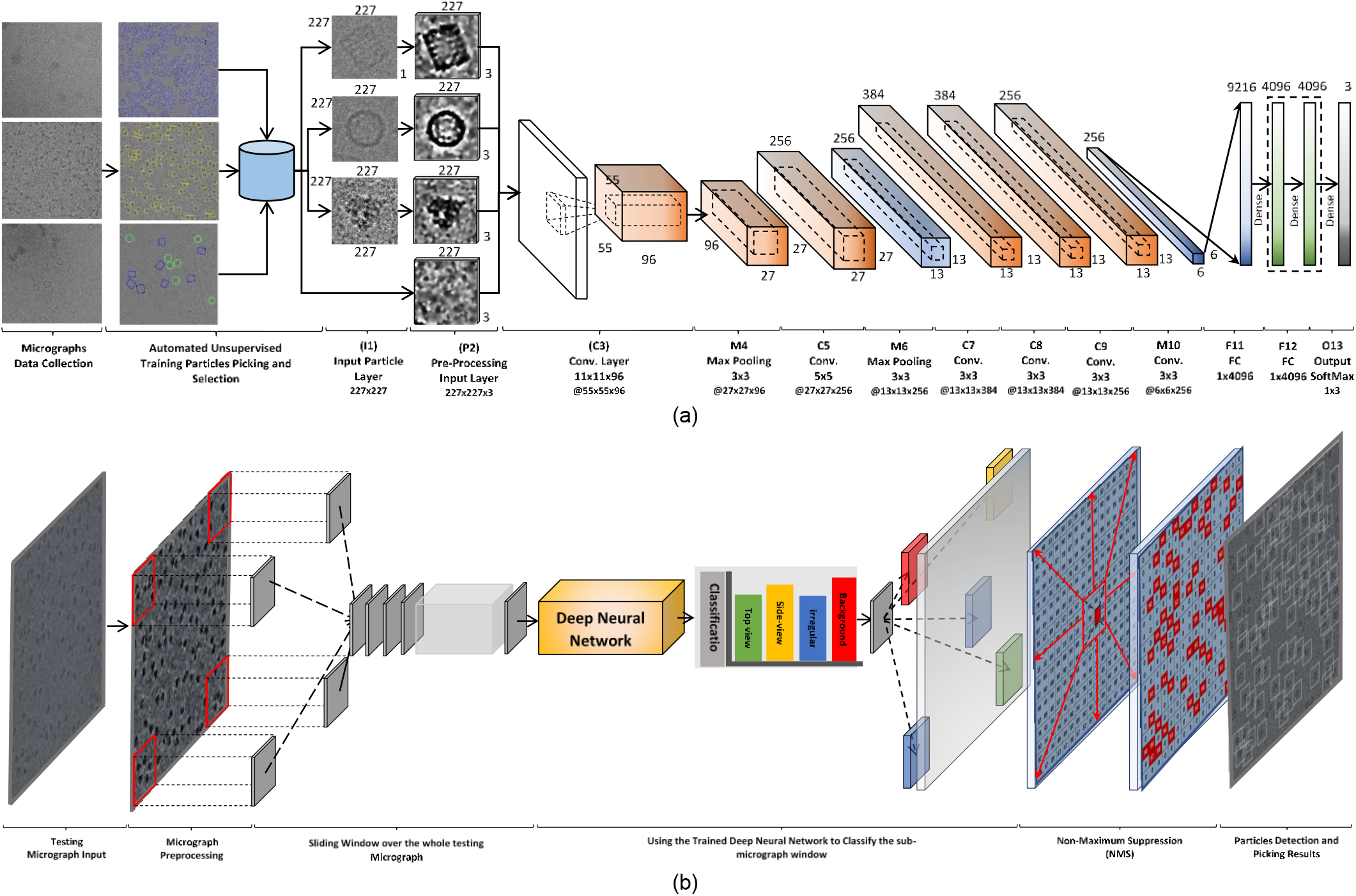
The architecture of the deep neural network used in DeepCryoPicker. (a) Training pipeline, (b) Testing pipeline. The convolutional layer and the subsampling layer are abbreviated as C and S, respectively. C3:11×11×96 means that in the third convolutional layer (C3) is comprised of 96 feature maps, each of which has a size of 11 × 11, also. C3: @27×27 means that output feature maps dimensions are 27×27 pixels.

**Table 5.** DeepCryoPicker network architecture

| Layer | Type | Filters | size |
| --- | --- | --- | --- |
| I1 | Input Layer | - | $227 \times 227 \times 3$ |
| P2 | Pre-processing | - | $227 \times 227 \times 3$ |
| C3 | Convolution | 96 | $11 \times 11$ |
| M4 | Max-pooling | - | $3 \times 3$ |
| C5 | Convolution | 256 | $5 \times 5$ |
| M6 | Max-pooling | - | $3 \times 3$ |
| C7 | Convolution | 384 | $3 \times 3$ |
| C8 | Convolution | 384 | $3 \times 3$ |
| C9 | Convolution | 256 | $3 \times 3$ |
| M10 | Max-pooling | - | $3 \times 3$ |
| F11 | Fully Connected | - | $1 \times 4096$ |
| F12 | Fully Connected | - | $1 \times 4096$ |
| O13 | Output | - | $1 \times 1$ |

The convolutional and sub-sampling layers, which are core building blocks of the convolutional neural networks (CNN), produce feature maps. The kernel sizes are selected to establish the local connections while expanding through the entire particle image. The learnable kernels are convolved with each feature map from the previous layer. The convolutional layers (in the same convolutional operations) share the same local connective weights 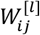 based on the previous layer’s weights 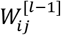, in which the feature maps in the current layer 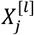 are produced based on Equation (3) [33]:

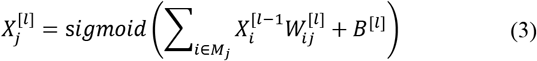

where *l* represents the convolutional layer, *W* and *B* is the shared weights and bias, *M* is extracted feature maps (in the previous layer), *j* is the output feature maps. Then, the feature maps are transformed to another layer by a non-linear activation function (sigmoid) as is given in Equation (3) [5]:

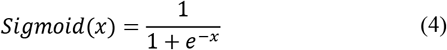

To reduce the positional over-fitting, the subsampling (max-pooling) layer is designed to subsample the input feature maps by decreasing the actual size and reduce the number of the parameters [33]: The max-pooling (subsampling) after each particular layer is computed based on Equation (5) [33]:

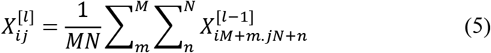

where *I* and *j* are the position of the output feature maps, *M* and *N* are the subsampling size. In the training process, the weights and bias are randomly initialized [0-1]. Then, they are updated during the training process. In our model, we used the cross-entropy loss function as the objective function Equation (6) [34]:

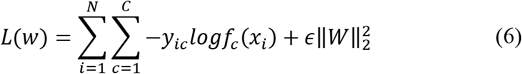

where *i* is the sample number and c is its label, *x* represents the predicted probability of the class *c*. *N* is the total number of training samples, and *C* is the total number of classes. During the training process, the errors of the objective function is minimized propagating error via the backpropagation algorithm based stochastic gradient descent as follow [35] [36] [37].

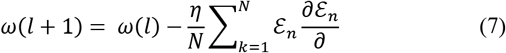

where *ɛ* is calculated as follow :

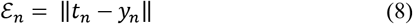

where *t*_*n*_is the label of the *n*^*th*^ training sample, and *y*_*n*_ is the value of the output layer corresponding to the *n*^*th*^ training sample. *ω*(*l*) and *ω*(*l* + 1) represents the training parameter before and after the update of each iteration. The learning rate,*η*, is initially set to 0.0001.

### Performance evaluation metrics

For the evaluation of the performance results we use one of the most popular evaluation metrics which is the precision-recall curve [34] that are defined by Equation (9) and (10), accuracy and F1 score that are defined by Equations (11) and (12) respectively [34].

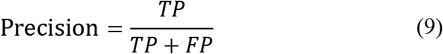

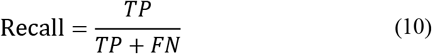

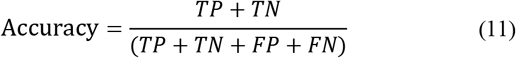

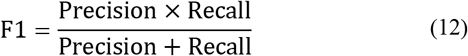

where TP is true positives of particles that are correctly picked among the total particles number, FP is the false positives of other objects that are incorrectly detected as particles. FN (false negatives) is particles that are incorrectly predicted as non-particles.

### Data augmentation

During the training, instead of passing the original particle image, each image is augmented by using the preprocessed version of the same particle image before passing it through the deep network.

## Data and code availability

The datasets used in this study and the source code of DeepCryoEM are available at https://github.com/jianlin-cheng/DeepCryoEM.

## Acknowledgements

Research reported in this publication was supported in part by two NSF grants (DBI 1759934 and IIS1763246) to JC, an NIH grant (R01GM093123) to JC and JT, and an administrative supplement to R01GM065546 (Collaborative Supplements for Cryo-Electron Microscopy Technology Transfer) to JT.

## Author contributions

JC conceived of the project. AA and JC designed the experiment. AA implemented the method and gathered the results. AA, JC, AO and JT analyzed the data. AA and JC wrote the manuscript. All authors edited and approved the manuscript. The manuscript is written by AA and has been proofread by MH and JC. AA, JC and YD discussed the results, new method and the manuscript.

## Additional information

**Supplementary Information**: some additional figures and algorithms are provided in the Supplementary File.

